# Metannot: A succinct data structure for compression of colors in dynamic de Bruijn graphs

**DOI:** 10.1101/236711

**Authors:** Harun Mustafa, André Kahles, Mikhail Karasikov, Gunnar Rätsch

## Abstract

Much of the DNA and RNA sequencing data available is in the form of high-throughput sequencing (HTS) reads and is currently unindexed by established sequence search databases. Recent succinct data structures for indexing both reference sequences and HTS data, along with associated metadata, have been based on either hashing or graph models, but many of these structures are static in nature, and thus, not well-suited as backends for dynamic databases.

We propose a parallel construction method for and novel application of the *wavelet trie* as a dynamic data structure for compressing and indexing graph metadata. By developing an algorithm for merging wavelet tries, we are able to construct large tries in parallel by merging smaller tries constructed concurrently from batches of data.

When compared against general compression algorithms and those developed specifically for graph colors (VARI and Rainbowfish), our method achieves compression ratios superior to gzip and VARI, converging to compression ratios of 6.5% to 2% on data sets constructed from over 600 virus genomes.

While marginally worse than compression by bzip2 or Rainbowfish, this structure allows for both fast extension and query. We also found that additionally encoding graph topology metadata improved compression ratios, particularly on data sets consisting of several mutually-exclusive reference genomes.

It was also observed that the compression ratio of wavelet tries grew sublinearly with the density of the annotation matrices.

This work is a significant step towards implementing a dynamic data structure for indexing large annotated sequence data sets that supports fast query and update operations. At the time of writing, no established standard tool has filled this niche.

## 1 Introduction

### 1.1 Background

The ever-decreasing cost of high-throughput sequencing (HTS) has led to massive growth in the availability of DNA and RNA sequencing data to researchers and the greater scientific community [21]. Several large-scale projects, such as the 1000 Genomes Project [4], UK10K [33], and many others [32, 35], have enabled us to much more extensively sample the genetic variation among humans and other organisms of interest. In addition to providing raw sequencing reads, follow-up projects such as ExAC and gnomAD have consolidated some of this data into large variant call sets to facilitate subsequent analysis [16]. However, the rate of sequencing data generation continues to exceed the rate at which these data can be indexed, processed, and analyzed [21].

Traditional sequence indexing and search methods, such as hash table-based seed-and-extend [2] or Burrows-Wheeler transform (BWT)-based [6] read-to-reference alignment [18], are optimized for relatively small^1^ databases of long reference sequences, and thus, HTS data have remained largely unindexed and not efficiently searchable. The development of metagenomics has complicated the issue even further, since sequence information from millions of as-of-yet uncharacterized organisms is available only in HTS data sets [32, 9]. Without sufficient sequencing data or reference genomes to properly assemble the genomes of individual species from these samples, much of this valuable data is currently difficult to process for specialists and inaccessible to non-specialists from the greater research community.

### 1.2 Recent models for metagenome indexing

Recent models for sequence indexing can be divided into two main groups: *hashing-based* and *graph-based*.

Hashing-based methods use probabilistic data structures for lossy or lossless compression of sequences, graph elements, or metadata, and allow for fast approximation of various queries, such as similarity between pairs of sequences [23], membership in a set of sequences [24], or subsequence counting [36]. Recently, sequence Bloom trees [29] and split-sequence Bloom trees [30] have been introduced for indexing HTS data. However, due to their use of Bloom filters for sequence matching, they require recomputation with different parameters as they saturate to maintain a given false positive rate. Each hashing method is typically optimized for performing a narrow range of queries, and thus, a separate copy must be stored for every query type supported.

Graph-based methods were first used for assembling short read sequencing data into long contiguous sequences (contigs) [27]. Most of these can be described as variants of de Bruijn graphs [34], overlap graphs [22], cactus graphs [25], and others [26]. The succinct representation of the uncompacted de Bruijn graph by Bowe, Onodera, Sadakane, and Shibuya [5] (henceforth referred to as BOSS) has acted as the basis for sequencing projects where the sheer sizes of the input data, such as metagenomics data sets, have necessitated trading increased running times for dramatically decreased storage [15, 17, 24, 20].

In order to use a de Bruijn graph as the backend for a sequence search method, however, an additional method must be developed to encode and compress associated metadata (which we refer to as *graph colorings*). When query sequences are mapped to paths on the graph, these paths induce sequences of colors annotating sections along the paths. Colors on edges can be used as indicators for various metadata categories (given some ordering of categories), such as their presence in certain samples, genetic structures, or their implications in diseases[20]. These are encoded as a large bit matrix (which we refer to as the *annotation matrix*), with one row for each edge and one column for each metadata category. One of the early methods for color encoding is the positional BWT [8], where sample haplotypes are encoded as bit vectors on a reference sequence and a BWT is applied before they are compressed. This method has also been extended to work with positions on string graphs [22].

Recent methods using the BOSS representation for colored de Bruijn graphs, such as Bloom filter tries [14], VARI [20], Rainbowfish [1], and deBGR [24], have proposed methods for succinct compression of graph colors. The VARI pipeline concatenates the rows of its bit matrix and compresses by Elias-Fano [11, 10] or Raman-Raman-Rao (RRR) [28] coding depending on the proportion of set bits [20]. The method in Rainbowfish builds on this by computing Huffman codes for the edge colors and compressing the concatenations of the codes by RRR coding [1]. Bloom filter tries are a probabilistic data structure for storing the edge labels and colors of a colored de Bruijn graph [14], while deBGR encodes these in a quotient filter and uses the colors of neighboring edges for error correction [24]. Although Rainbowfish achieves the best compression ratios among the lossless methods, its use of Huffman codes does not take full advantage of correlations between columns in the annotation matrix. In addition, it requires the distribution of the edge color frequencies to be known beforehand.

These methods rely on static data structures for optimal compression, requiring full decompression, extension, and recompression steps to perform edits. In the case of Rainbowfish, extensions can potentially reduce the efficiency of the coding if novel colors follow a different distribution from those which were used to compute the codes. In the worst case, a full recomputation of all Huffman codes would be required at regular intervals to maintain a desired compression efficiency. The static natures of these methods renders them inadequate for use as backends in dynamic sequence databases.

We have recently been developing methods for the fast construction and storage of the BOSS representation of de Bruijn graphs in both static (for fast querying) and dynamic (for fast updates) data structures, with the ability to convert between internal representations depending on the desired types of user interaction. In this work, we further extend these methods by proposing a dynamic data structure for graph colorings compression that takes advantage of correlations between columns of the annotation matrix and can be combined with other models for sequence indexing as well.

### 1.3 Wavelet tries: a dynamic data structure for annotation compression

For the compression of graph colorings, we propose a novel application of the *wavelet trie* data structure [13]. Briefly, a wavelet trie is an extension of the concept of a wavelet tree and takes the shape of a compact prefix tree (a binary radix trie). Instead of compressing strings over a fixed alphabet, wavelet tries compress tuples of bit vectors, where each vector is the binary encoding of a string over an alphabet of arbitrary size. This allows the structure to compress dynamic strings over arbitrary alphabets by finding common contiguous subsequences (or *segments*) among the bit vector encodings of its characters. In the context of genome graph coloring, the tuple of bit vectors can be defined as the rows of the annotation matrix in which the number of columns grows to encode new categories of metadata. In the worst case, the height of a wavelet trie is equal to the length of the longest bit vector being compressed (in the case where no common prefices are present at every internal node).

To the authors’ knowledge, no implementation of this data structure has been reported. In this manuscript, we present an implementation employing a parallel construction strategy via wavelet trie merging. The merging algorithm presented is a generalization of the algorithm provided by the original manuscript for appending novel bit vectors to an existing wavelet trie [13].

## 2 Methods

### 2.1 Succinct de Bruijn graph construction

The Bowe, Onodera, Sadakane, and Shibuya (BOSS) representation of the de Bruijn graph was chosen as the underlying genome model for this study [5]. Let *k* be some fixed positive integer and *G* be a de Bruijn graph of order *k*. When the edges of *G* are sorted by the reverse lexicographical ordering of the *k*-mer labels of their respective source nodes (using their own edge labels as tie breakers), only the last character of each *k*-mer (vector *F*), the edge labels (vector *W*), and two auxiliary bit vectors (vectors *ℓ* and *W*^−^) need to be stored to represent the graph. *ℓ* is an indicator for the last outgoing edge of a node, while *W*^−^ is an indicator for all but the first edge leading to a node with an in-degree greater than one. In this representation, there is a one-to-one correspondence between *F* and *W*, where the ith occurrence of a character *c* in *W* with *W*^−^ value 0 corresponds to the *i* occurrence of *c* in *F* with *ℓ* value 1 [5]. Construction of the BOSS representation of de Bruijn graphs is done using a binned parallel approach [5, 17].

### 2.2 Graph coloring during de Bruijn graph construction

Colors are computed for each edge of the de Bruijn graph during construction based on the metadata of the input sequences from which they are derived. During *k*-mer enumeration, assign each unique metadata category a positive integer ID and use these IDs to assign each *k*-mer a list of category IDs corresponding to its associated metadata categories. Then, convert the list of IDs to a bit vector (called an *edge color*) such that the IDs determine which bits in the vector are set to 1. When duplicate edges are removed during graph construction, combine their respective bit vectors via bitwise OR operations to define the new color of the remaining edge. Alongside the succinct de Bruijn graph, this process results in an auxiliary *annotation matrix* with n rows corresponding to the edges of the graph and *m* columns corresponding to the total number of unique metadata strings observed during construction. The resulting graph-matrix pair is a *colored de Bruijn graph*. When this graph is queried, sequences are mapped to a path (a sequence of edges) and a corresponding sequence of annotation matrix rows.

### 2.3 Graph color compression with wavelet tries

To greatly reduce the required storage space of the annotation matrix, while allowing for dynamic extension and random access to matrix rows, we chose to employ the *wavelet trie* data structure.

**Definitions and notation** Given a bit vector *b* ∈ {0,1}* (a finite string over the binary alphabet {0,1}), we use the notation |*b*| to refer to its length, *b*[*i*] to refer to its *i*th character, 1 ≤ *i* ≤ |*b*|, *b*[*j : k*] to refer to the bit vector *b*[*j*] ⋯ *b*[*k*], *b*[: *k*] to refer to its prefix *b*[1 : *k*], and *b*[*j* :] to refer to its suffix *b*[*j*] ⋯ *b*[|*b*|]. The empty vector is denoted *ε*.

The function **rank_0_**(*b, j*) (**rank_1_**(*b, j*)) counts the number of 0(1) characters in *b*[: *j*], while **select**_0_(*b, j*) (**select**_1_(*b, j*)) returns the index of the *j*^th^ 0(1) in *b*. Also, we will use the notation 2^*A*^ to denote the power set of a set *A* and abuse the notation |·| to refer to both set cardinalities and bit vector lengths.

**Construction** The wavelet trie encoding the annotation matrix *A* ∈ {0,1}^*n×m*^ is constructed recursively and is a binary tree of the form *T* = (*V,E*) (see Figure 1), where its nodes *n_j_* ∈ *V, j* ∈ {1,…, |*V*|} are of the form

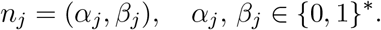

**Fig. 1:**
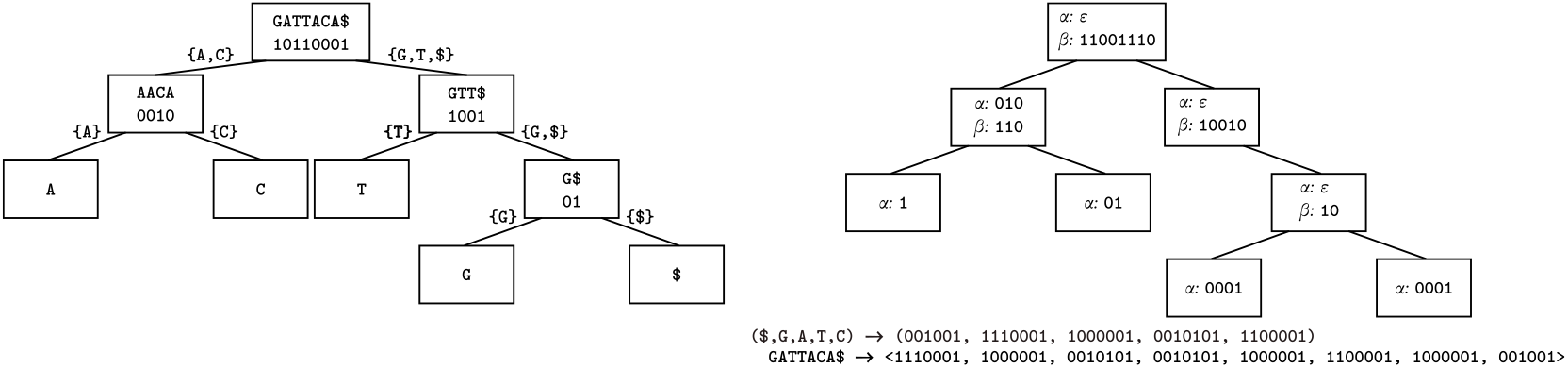
A wavelet tree (left) and a wavelet trie (right) constructed for the string GATTACA$ and the binary encodings of its characters, respectively. Wavelet tree. Characters are each node of the wavelet trie are divided equally amongst its two children, with a bit vector indicating these assignments. A node is a leaf when it is assigned only one character. In internal nodes, only the bit vectors need to be stored to allow traversal to the correct leaf when querying the wavelet tree. **Wavelet trie**: In a wavelet trie, strings are encoded as tuples of bit vectors. At a node, the common prefix of the bit vectors is extracted and the next significant bit is used to assign the bit vector suffices to that node’s children. A node is a leaf when all bit vectors assigned to it are equal. In both structures, index queries are resolved by traversing the tree and performing rank operations on the distribution bit vectors. In this example, ASCII codes are used to define the binary codes for each character.

The *α_j_* are referred to as the *longest common prefices* (LCPs) and the *β_j_* are referred to as the *assignment vectors*.

The algorithm starts with the *root* node *n*_1_. We define the initial set of input bit vectors to be the rows of *A*, 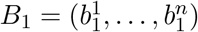, where 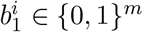 for all *i* ∈ {1,…, *n*}.

On the *j*th iteration, for a list of input bit vectors 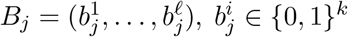, 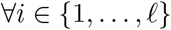, ∀*i* ∈ {1,…,*ℓ*}, compute *n_j_* as follows: Compute the longest common prefix 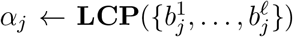 for the bit vectors in *B_j_*. Formally, this function is defined as follows, **LCP** : 2^{0,1}*^ → {0,1}*,

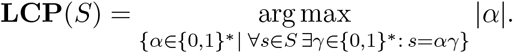

If the computed *α_j_* matches all the input bit vectors, *n_j_* is referred to as a *leaf* and let the assignment vector be *β_j_* ← *ε*. Then terminate this branch. Otherwise, the set the assignment vector to be the concatenation of next significant bits in each of the 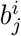, *i* ∈ {1,…, *ℓ*} after removing the common prefix *α_j_*,

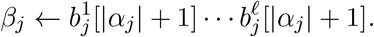

Continue the recursion on the child nodes *n*_2*j*_ and *n*_2*j*+1_ with the new sets of bit vectors *B*_2*j*_ and *B*_2*j*+1_, respectively, which are defined by partitioning *B_j_* based on *β_j_* and removing the first |*α_j_*|+2 bits,

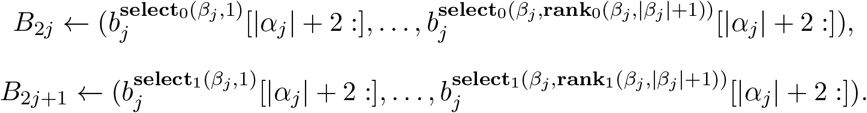

### 2.4 Parallel construction via wavelet trie merging

To allow for parallel construction of wavelet tries, we developed an algorithm to merge wavelet tries as a generalization of the wavelet trie extension method [13]. Merging proceeds by performing an *align* and a *merge* step on each node, starting from the root (see Figure 2 for an illustration of the process). Given two wavelet tries *T*′ and *T*″ with node sets 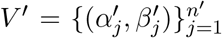 and 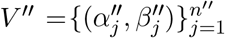 that we want to merge into a new trie *T*, the merging process can be summarized as:

1. **Align**: for the nodes 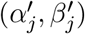 and 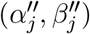, compute the longest common prefix 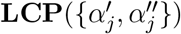 and make new nodes with this value and appropriate *β* vectors, set this to be the parent of the current nodes,
2. **Merge**: once 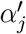 and 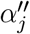 are equal, concatenate 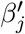 and 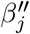,
3. **Repeat**: move down to *j*’s children and apply the same function until all leaves are reached.

**Fig. 2:**
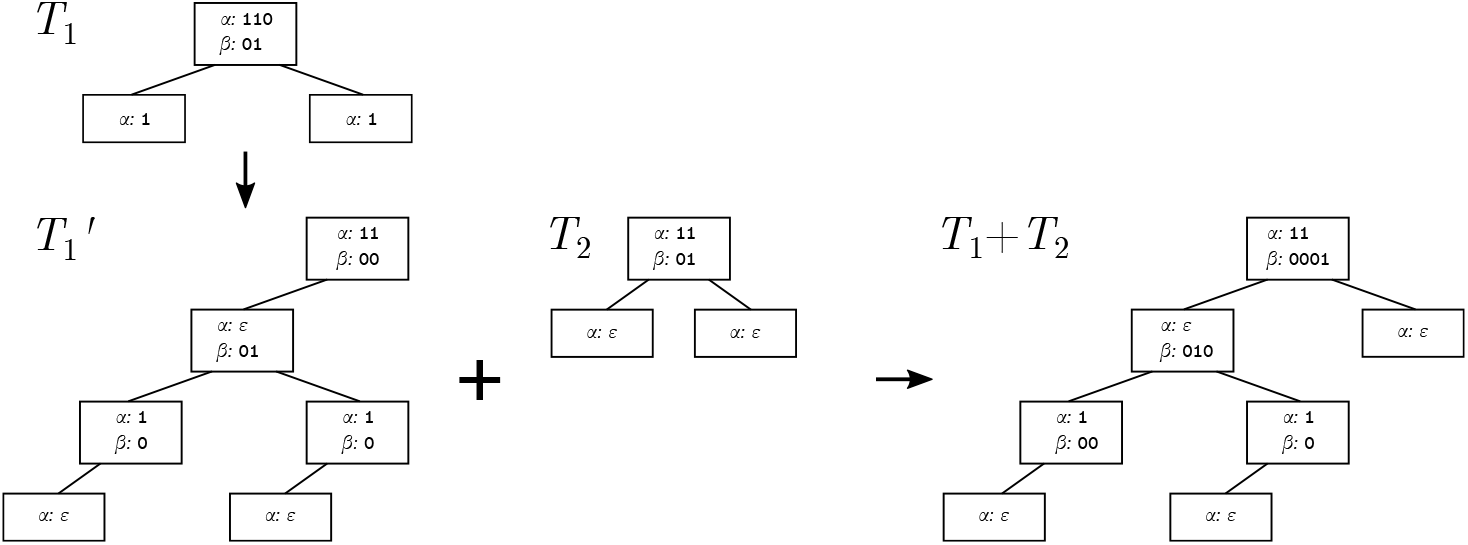
Merging of wavelet tries *T*_1_ and *T*_2_ to form the wavelet trie *T*_1_ + *T*_2_. Starting from the root node, the common prefix of the two *α* vectors is found and new *β* vectors are computed from the their remainders. These become new parent nodes and the initial nodes’ *α* vectors are updated to their respective remainders after removing the common prefix (e.g., the conversion from from *T*_1_ to 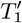). When the two *α*s are equal, their respective *β*s are concatenated and the merging function is applied to their children. When a leaf is reached in one tree, but the equivalent node in the other tree is internal (e.g., the left child of the root in *T*_1_ + *T*_2_), the leaf is merged by appending or prepending additional zeros to the *β* vectors of all left ancestors. Note that extra leaf nodes producing trailing zeros in the decoded bit vectors are added during the merging process. See Section 2.4 for more details.

In the context of compressing the edge colors of a de Bruijn graph, this method assumes that the columns of two wavelet tries being merged are indicators for matching metadata categories.

For this method, we define the *descendants* function **D** : {1,…, |*V*|} → 2^{1,…,|*V*|}^ for the wavelet trie *T* = (*V, E*) with nodes 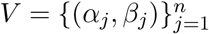 by the recurrence

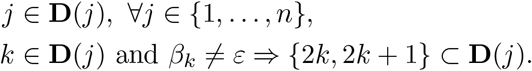

The three steps in the merging operations are as follows:

**Align** Given nodes 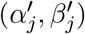 and 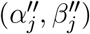, we compute their longest common prefix

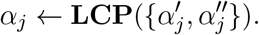

If 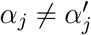, we let

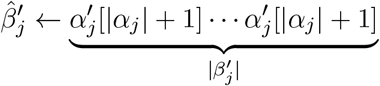

and update the indices in *T*′ by applying the transformation 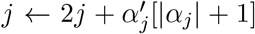 and updating all nodes *k* ∈ **D**(*j*) accordingly. We then let 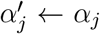 and 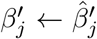 and truncate the prefix in the newly created child nodes,

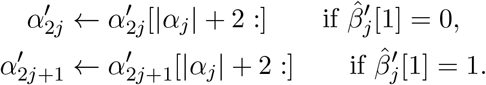

If 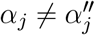, the second trie is processed accordingly.

**Merge** If 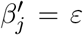 and 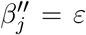, then terminate. Otherwise, if 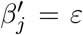, set 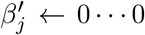 (of length 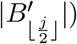). (similarly for 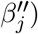). Then, merge the two assignment vectors

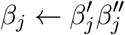

**Repeat** The merging algorithm is then performed on nodes *n*_2*j*_ and *n*_2*j*+1_ depth-first to continue the recursion.

If two wavelet tries constructed from bit vectors of different lengths are merged, this merging algorithm leads to the decoding of bit vectors with trailing zeros. Since we indend to use these vectors as indicators for various metadata, the presence of extra trailing zeros in the decoded bit vector does not represent false information.

### 2.5 Computational complexity of wavelet trie operations

Let *A* ∈ {0,1}^*n×m*^ be an annotation matrix. The height of a constructed wavelet trie *T* = (*V, E*) depends on the degree to which the bit vectors share common segments, with the worst-case value being *h* ≤ min(*n, m*) when no segments are shared. Since there can be at most *n* leaves, and the maximum height of the tree is at most *m*, the number of nodes can be at most |*V*| ≤ min(*n*, 2^*m*^).

Given two wavelet tries *T*_1_ and *T*_2_, merging is performed in 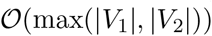 time. Once a tree is constructed, queries can be performed in 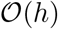 time. To achieve this value, the *β_j_* are compressed with RRR coding [28] to support rank operations in 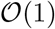 time.

### 2.6 Improving compression ratios using graph topology

One of the advantages of maintaining a graph-based model for genome storage is its ability to efficiently represent an ordering on the *k*-mers. On the other hand, the ordering provided by the BOSS representation is rarely optimal for compressing adjacent edge colors by run-length encoding, since adjacent edges frequently come from different samples. There is, however, an additional degree of freedom in that the ordering of annotation matrix columns is not fixed. Since the compression ratio of a wavelet trie depends on both the ordering of the rows (defining the compressability of the *β*s) and the similarity of the bit vectors after each assignment during construction (which defines the height and balance of the tree), we explored whether graph structure could be used to help provide additional prefix bits to help optimize row segregation.

If we assume that a certain set of paths in the de Bruijn graph (i.e., those corresponding to reference genomes) act as *backbones* (whose indicator columns in the annotation matrix we refer to as *backbone bits*), while other paths represent sequence variation, then it is expected that the edge colors of backbone paths are highly correlated with those of the variation paths. The edge colors of variation paths can be supplemented by setting the columns of their corresponding backbones to 1.

We now describe this process more precisely. Let *b*^1^,…, *b^n^* be the rows of the input annotation matrix and let *C* = {1,…,*m*}, where *m* is the number of columns of the annotation matrix. Let *R* ⊂ *C* be the set of indices/IDs generated from backbone paths/genomes and let *P* map elements of *R* to their corresponding paths in *G*. The user provides a map **B** : *C* → *C* such that *R* = {*j* |*B*(*j*) = *j j* ∈ *C*} is the set of fixed points of **B**. Then, for each *j* ∈ *C* and each *b^i^* s.t. *b^i^*[*j*] = 1, we set *b^i^*[**B**(*j*)] ← 1.

When this process is not followed, we say that the backbone bits are *unset*, whereas applying this process results in the backbone bits being *set*. For example, given an index *i* corresponding to a backbone and *j* corresponding to a variant, we say that the backbone bit is set if *i* = *j* = 1 and unset if *i* = 1 and *j* = 0.

## 3 Results

The following section covers our evaluation of the wavelet trie data structure on a variety of data sets. This includes a comparison of its compression ratio against general compression algorithms and to those developed specifically for graph colors. In addition, we evaluate the hypothesis that setting backbone bits using prior knowledge improves compression ratios. Finally, we study how the compression ratio of wavelet tries behaves as a function of the number of metadata categories m and the density of the annotation matrix (the ratio of the number of bits set to 1 and nm) a linear hierarchy (called a *chain*) of models ranging from 50 to 1000 virus genomes.

### 3.1 Data sets

Data sets originating from viruses (Virus100 and Virus1000), bacteria (simply Bacteria), and humans (chr22+gnomAD and hg19+gnomAD) are used in this study to construct graphs with varying topologies to study their effects on the wavelet trie’s compression ratios. See Appendix Section A.1 for a precise description of the data sets used.

### 3.2 Wavelet trie compression ratios similar to gzip and bzip2, and better than previous methods

As baseline comparisons, the compression ratio of wavelet tries was compared to those of the standard UNIX compression utilities gzip and bzip2 (see Table 1). gzip is an implementation of the LZ77 algorithm and encodes blocks of text, while bzip2 performs a sequence of transformations, including run-length encoding, BWT, move-to-front transforms, and Huffman coding. In addition, the compression performance of wavelet tries was compared to other methods developed specifically for annotation matrices on succinct de Bruijn graphs, such as the methods presented in VARI [20] and Rainbowfish [1]. We measured *compression performance* as numbers of bits stored by the structures (which we denote *s*) divided by the total number of bits in the matrices (*nm*).

**Table 1:** Compression performance of wavelet tries and other algorithms. Performance is measured as the average number of bits per edge. Wavelet tries on all but the Virus1000 data set outperformed gzip and the VARI method, but were outperformed by bzip2 and the Rainbowfish method. At the time of writing, VARI was unable to encode the Virus100, Virus1000, and hg19+gnomAD data sets. The quantities s and nm refer to the number of set bits and the total number of bits in the annotation matrices, respectively, with backbone bits unset. Thus, the quantity in the third column is a measure of matrix density. The headings **WTr, Wtr** (**bb**), and **RF** refer to “wavelet trie”, “wavelet trie with backbone bits set”, and “Rainbowfish”, respectively.

| Data set | # edges/$10^6$ | $\frac{s}{nm}$ (%) | bits/edge | WTr | WTr (bb) | gzip | bzip2 | VARI [20] | RF [1] |
| --- | --- | --- | --- | --- | --- | --- | --- | --- | --- |
| Virus100 | 2.052 | 2.002 | 100 | 8.025 | 8.196 | 15.576 | 6.902 | N/A | 15.090 |
| Virus1000 | 15.360 | 0.216 | 1000 | 67.174 | 22.756 | 38.239 | 12.606 | N/A | 36.011 |
| Bacteria | 18.713 | 7.936 | 45 | 6.979 | 5.876 | 8.805 | 5.311 | 33.761 | 4.851 |
| chr22+gnomAD | 54.723 | 1.703 | 10 | 3.221 | 3.159 | 5.860 | 2.990 | 18.498 | 2.464 |
| hg19+gnomAD | 2886.802 | 3.287 | 31 | 7.510 | 7.740 | 9.345 | 5.291 | N/A | 5.573 |

The results indicate that wavelet trie compression outperforms gzip and the VARI method. bzip2 and the Rainbowfish method achieve similar compression ratios and slightly outperform our method. The Virus1000 data set is notable in that wavelet tries exhibit the worst compression performance among the methods tested, though much better results were achieved when backbone bits were set. At the time of writing, the VARI method was unable to compress the annotations for the Virus100, Virus1000, and hg19 data sets. Setting backbone bits led to a three-fold improvement in the compression performance on the Virus1000 data set (from 67.174 bits per edge to 22.756), marginal improvements in the compression performance on the bacterial and chr22 data sets, and a marginal decrease in performance on the Virus100 and hg19 data sets.

### 3.3 Setting backbone bits improves compression ratios

To test the hypothesis that the setting backbone bits (which by definition tend to occur in columns with lower indices) reduces compression ratios, 100 random shufflings of the column ordering in the Bacteria and Virus100 data sets were generated and the resulting data compressed to approximate the null distribution of compression ratios (see Figure 3). The results indicate that the original ordering of columns was optimal with respect to the defined null distribution when backbone bits were set. As a negative control, when the backbone bits were unset, the resulting compressed file sizes did not significantly differ from the means of the null distributions.

**Fig. 3:**
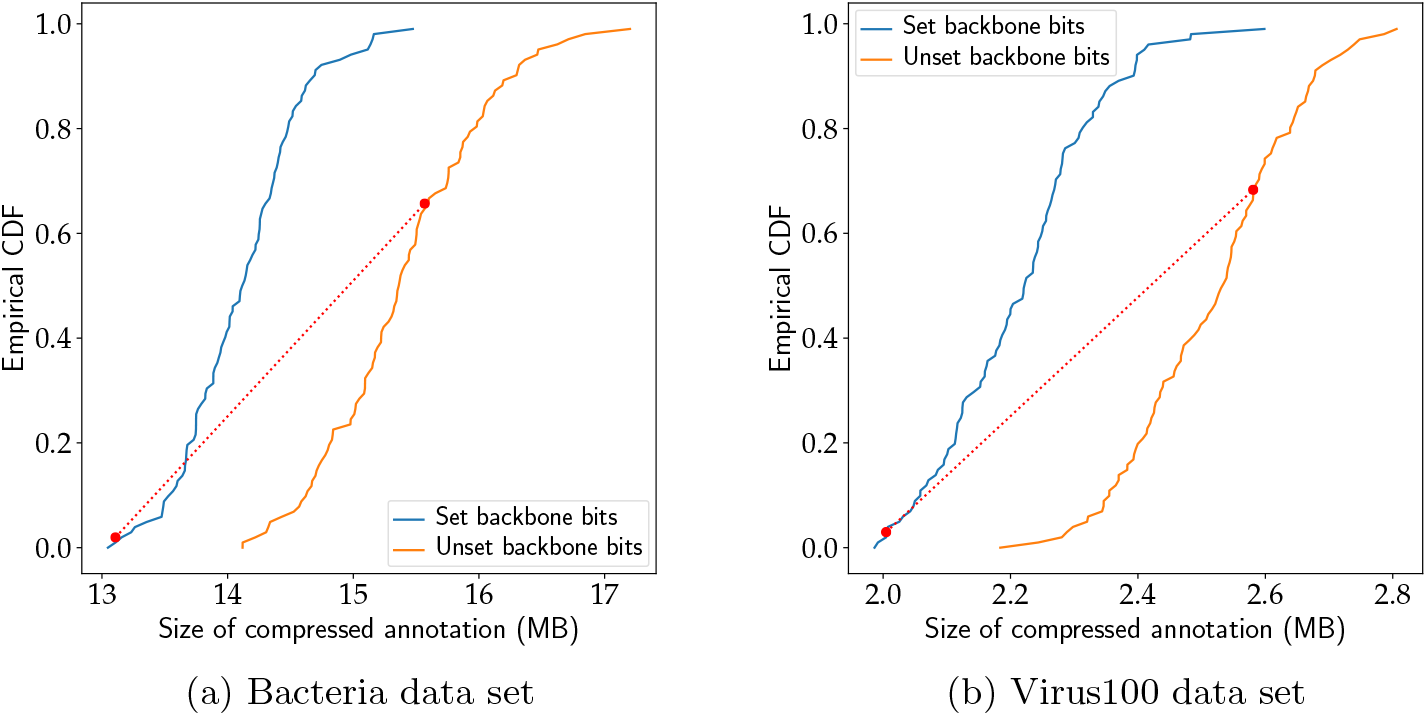
Distribution of the file sizes of wavelet tries over 100 random permutations of the annotation matrix column order. The red line indicates the mapping between the positions in the CDFs of the input column ordering determined by the graph. Setting the backbone bits in both cases leads to a decrease in the sizes of the compressed files. On both data sets, the originally ordering of the columns leads to an optimal compression ratio when backbone bits are set, an unoptimal ratio otherwise.

### 3.4 Wavelet trie size grows linearly with increased unique compression size

To test the scalability of wavelet trie compression, we generated a *chain* (a linear hierarchy) of virus graphs ranging from 50 to 1000 random genomes in steps of 50 (i.e., *G*_1_ ⊂ ⋯ ⊂ *G*_20_) and measured the compression ratios of the annotations for each graph. The compression ratios for the Virus50 to Virus1000 graphs exhibit an exponential drop from 12.5% to 6.5% when backbone bits are not set and a steeper drop from 12% to 2% when backbone bits are set (see Figure 4). The compression ratios grew sublinearly as the density of the annotation matrices grew, with little difference in the growth characteristics with and without the backbone bits set. By definition, the matrix densities tended to be marginally greater when backbone bits were set.

**Fig. 4:**
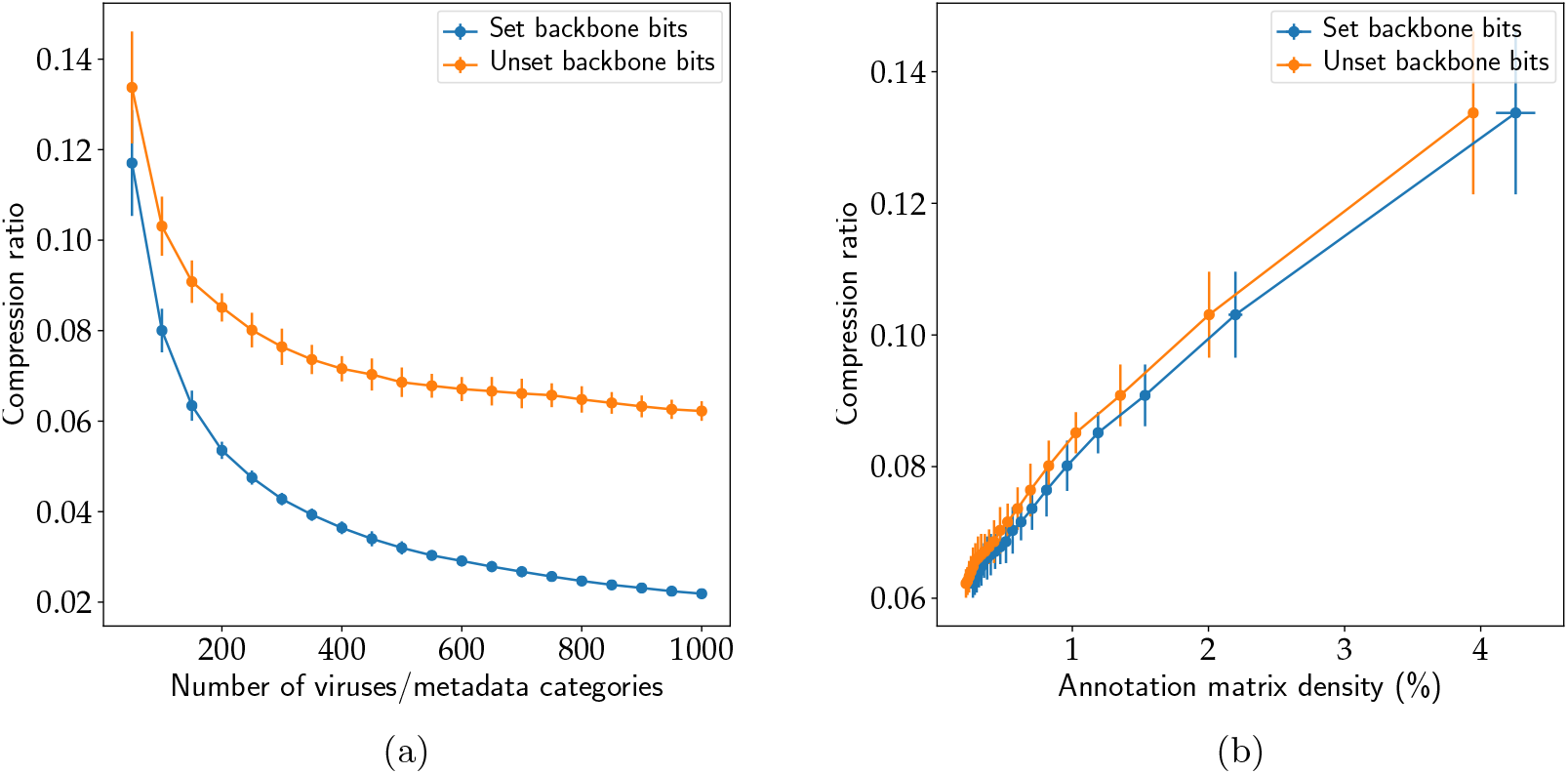
Compression ratio of the wavelet trie on virus graphs of increasing (a) genome count and (b) annotation matrix density. An exponential drop in compression ratio, converging at 6.5%, is observed as the number of metadata categories increases, with a steeper decline converging at 2% when backbone bits are set. Analogously, a sublinear increase in the compression ratio was observed as the density of the annotation matrix increased. The compression ratio is defined as the ratio of the file size of the serialized wavelet tries and the total number of bits in the input annotation matrices (i.e., the number of edges multiplied by the number of virus species). The annotation matrix density is defined as the percentage of matrix bits which are set to 1. The Virus1000 graph was generated from a sampling of 1000 virus genomes from GenBack (see Section 3.1), while the other 19 graphs were generated by taking the first *j* genomes in Virus1000, where *j* ranged from 50 to 1000 in steps of 50. The trials were repeated 10 times with different random seeds, with the curve and error bars representing the mean and standard deviations of the compression ratios, respectively.

## 4 Discussion and Conclusions

In this study, we have tacked the problem of encoding sequence metadata as edge colors to construct succinct colored de Bruijn graphs. Given a binary matrix encoding of metadata (with one row per edge and one column per metadata category), we have presented a parallel construction method and novel application of the wavelet trie data structure for matrix compression. The construction method builds smaller wavelet tries on batches of data and merges them to form the full trie, performing every step in a multithreaded fashion. The resulting structure is dynamic in that novel edge colors of arbitrary size can be appended. In addition, we have demonstrated that when using indicators for the backbone regions of the de Bruijn graph positioned in low-index columns of the annotation matrix, we are able to improve compression ratios by assisting edge color segregation during wavelet trie construction. Thus, we are able to take advantage of graph topology to improve compression performance.

The results on the Virus1000 set of graphs indicate that our implementation of wavelet tries, for sufficiently large graphs, stabilize at compression ratios of 6.5% and 2% when backbone bits are unset and set, respectively. The data structure is less efficient on smaller graphs due to the greater significance of the employed data structures’ overhead. The high variability among viral sequences led to each added batch of sequences, on average, adding a constant amount of information to the graph and its compressed annotation. With regard to their effect on trie structure, each new batch creates a split in a node close to the root, forming a large separate subtrie, and thus, reducing the chances of such splits occurring with each subsequent extension. While these graphs represent cases of relatively few sequences with modest metadata category counts, larger graphs, such as those constructed on larger collections of eukaryotic genomes will need to be constructed to further study the wavelet trie’s growth characteristics.

One significant limitation of wavelet tries is their reliance on shared segments (contiguous subsequences), especially in the first few columns of the annotation matrix, to effectively partition the rows for optimal compression. While this is partially addressed by setting backbone bits in the annotation matrix, a more principled approach with less user input will become necessary in future releases. This would involve an analysis of the de Bruijn graph topology to algorithmically determine paths to use for backbone bits.

An alternate approach for compressing graph colors which does not rely on this analysis would be to use a directed acyclic graph as a model, as is typically applied for compressing dictionaries [3]. With this structure, matching nodes in different branches are merged into single nodes to further reduce redundancy. However, the computational complexity of dynamically maintaining such a structure, and the added complexity of preserving the ordering of the annotation rows (to support row index queries) are challenges that must be addressed for such an approach to be appropriate for compressing graph colors.

With regard to the original stated purpose of developing an annotated sequence graph for indexing reference sequence and sequencing reads, an ideal implementation of such a database would (1) employ a dynamic data structure which (2) supports fast queries and updates, and is (3) optimal with regards to storage. At the time of writing, no database solution fulfilling all three criteria on this type of data has been published. One observation, however, is that in real world applications, database updates are much less frequent than queries. Thus, a dynamic data structure with slower query times may, on average, underperform compared to one involving a static data structure with fast query times that is updated periodically. We propose a solution in which the backend is able to switch between static and dynamic states efficiently. This way, the average time complexity of query and update operations is kept low through amortization.

## 5 Acknowledgements

We would like to thank Torsten Hoefler, and the Biomedical Informatics Group at ETH Zurich, in particular Amir Joudaki, Viktor Gal, and Gideon Dresdner for helpful discussions and criticism. This project was funded by the Swiss National Science Foundation (SNF) grant #407540-167331 “Scalable Genome Graph Data Structures for Metagenomics and Genome Annotation” as part of Swiss National Research Programme (NRP) 75 “Big Data”. The authors declare no conflicts of interest.

## A Appendix

## A.1 Data sets

### Bacteria

This data set is composed of 45 strains of bacteria in GenBank [7] from the *Lactobacillus* species *acidophilus, amylovorus, brevis, buchneri*, and *casei*. The columns in this graph’s annotation matrix indicate presence of an edge in each of the strains. Because of the low variability in the input sequences, they are represented as a graph with a predominantly linear topology and short variant paths (called *bubbles*). One genome from each of the species was chosen as a backbone path. The resulting graph had 18,669,398 unique *k*-mers, 18,713,013 edges, and 536 unique edge colors (i.e., bit combinations). See Appendix Section A.3 for a list of the bacterial strains used.

### Virus1000

This data set is composed of 1000 virus genomes randomly selected from GenBank, meant to study a graph whose topology is a series of almost mutually-exclusive loops with slight variation. The columns in this graph’s annotation matrix indicate presence of edges in each of the virus genomes. Similar to the Bacteria data set, the viruses were grouped by the first word of their names and the first species in each group was assigned as a backbone path. The resulting annotation bit matrix is very sparse and adjacent rows are either almost identical or almost mutually exclusive. This graph contains 15,342,369 unique *k*-mers, 15,360,442 edges, and 10,585 unique edge colors.

### Virus100

This is a subset of the Virus1000 set containing only 100 virus strains used to facilitate the permutation tests in Section 3.3. This graph contains 2,051,777 unique *k*-mers, 2,052,501 edges, and 284 unique edge colors.

### chr22+gnomAD

This graph consists of chromosome 22 from the `hg19` assembly of the human reference genome as the main reference backbone. To provide genetic variability, the set of exome variants from the gnomAD data set were incorporated into the graph [16]. This larger data set is meant to analyze the properties of the trie when the underlying graph is large, but with little variability. The columns in this graph’s annotation matrix are defined as indicators for its edges’ presence in 9 ethnic groups defined in the data set. The first column in the matrix is used to indicate edges which are present in the reference genome and serves as the backbone bit. The graph contains 54,386,415 unique *k*-mers, 54,723,569 edges, and 595 unique edge colors.

### hg19+gnomAD

This graph was constructed from the same data sets as the one described above, using data from the full human autosome. The same definition is used for the annotation matrix columns, with 9 columns being used to indicate edges observed in the defined ethnic groups and 22 prefix columns being used to indicate presence in the first 22 reference chromosomes as the backbone bits. This graph’s topology was designed to be analogous to the Virus1000 data set, but with 1000 × the number of rows and one-tenth of the number of annotation columns. It contains 2,880,005,212 unique *k*-mers, 2,886,801,846 edges, and 320,856 unique edge colors.

## A.2 Implementation and source code availability

All algorithms were implemented in C++ 14 using the `boost` (arbitrary precision integers), `htslib` (VCF parsing) [19], `sdsl-lite` (static succinct data structures) [12], and **libmaus2** (dynamic succinct data structures) [31] libraries. Wavelet tries are stored in memory in a fashion similar to linked lists, with `Node` objects containing pointers the objects that define their children. For serialization, this structure is packed into a `std::unordered_map` data structure mapping node indices to `Node` objects.

Our implementation is provided as a header-only library and a standalone executable at http://www.github.com/ratschlab/metannot.

## A.3 List of bacterial strains used

– *Lactobacillus acidophilus*

- 30SC (uid 63605)
- La 14 (uid 201479)
- NCFM (uid 57685)
– *Lactobacillus amylovorus*

- GRL 1112 (uid 61179)
- GRL1118 (uid 160233)
– *Lactobacillus brevis*

- ATCC 367 (uid 57989)
- KB290 (uid 195560)
– *Lactobacillus buchneri*

- NRRL B 30929 (uid 66205)
- uid 73657
– *Lactobacillus casei*

- ATCC 334 (uid 57985)
- BD II (uid 162119)
- BL23 (uid 59237)
- LC2W (uid 162121)
- LOCK919 (uid 210959)
- W56 (uid 178736)
- Zhang (uid50673)

## A.4 List of virus strains used

Due to their large number, the lists of virus strains used are made available in the previously-linked GitHub repository.

## Footnotes

1 in terms of the number of sequences

## References

1. Almodaresi, F., Pandey, P., Patro, R.: Rainbowfish: A Succinct Colored de Bruijn Graph Representation. bioRxiv (2017)

2. Altschul, S.F., Gish, W., Miller, W.: Basic local alignment search tool. Journal of molecular biology 215(3), 403–10 (1990)

3. Appel, A.W., Jacobson, G.J.: The World’s Fastest Scrabble Program. Communications of the ACM 31(5), 572–578 (1988)

4. Auton, A., Abecasis, G.R., Altshuler, D.M.: A global reference for human genetic variation. Nature 526(7571), 68–74 (2015)

5. Bowe, A., Onodera, T., Sadakane, K.: Succinct de Bruijn graphs. In: Lecture Notes in Computer Science (including subseries Lecture Notes in Artificial Intelligence and Lecture Notes in Bioinformatics), pp. 225–235. Springer, Berlin, Heidelberg (2012)

6. Burrows, M., Wheeler, D.J.: A block-sorting lossless data compression algorithm. Systems Research Research R(124), 24 (1994)

7. Clark, K., Karsch-Mizrachi, I., Lipman, D.J.: GenBank. Nucleic acids research 44(D1), D67–72 (2016)

8. Durbin, R.: Efficient haplotype matching and storage using the positional Burrows-Wheeler transform (PBWT). Bioinformatics (Oxford, England) 30(9), 1266–72 (2014)

9. Ehrilich, S.D., Consortium), (M.: MetaHIT: The Eurpoean Union Project on Metagenomics of the Human Intestional Tract. Metagenomics of the Human Body (2011)

10. Elias, P., Peter, Efficient Storage and Retrieval by Content and Address of Static Files. Journal of the ACM 21(2), 246–260 (1974)

11. Fano, R.: On the number of bits required to implement an associative memory. Massachusetts Institute of Technology Project MAC, Cambridge (1971)

12. Gog, S., Beller, T., Moffat, A.: From theory to practice: Plug and play with succinct data structures. In: Lecture Notes in Computer Science (including subseries Lecture Notes in Artificial Intelligence and Lecture Notes in Bioinformatics), pp. 326–337 (2014)

13. Grossi, R., Ottaviano, G.: The Wavelet Trie: Maintaining an Indexed Sequence of Strings in Compressed Space. (2012)

14. Holley, G., Wittler, R., Stoye, J.: Bloom Filter Trie: an alignment-free and reference-free data structure for pan-genome storage. Algorithms for Molecular Biology 11(1), 3 (2016)

15. Iqbal, Z., Caccamo, M., Turner, I.: De novo assembly and genotyping of variants using colored de Bruijn graphs. Nature Genetics 44(2), 226–232 (2012)

16. Lek, M., Karczewski, K.J., Minikel, E.V.: Analysis of protein-coding genetic variation in 60,706 humans. Nature 536(7616), 285–291 (2016)

17. Li, D., Liu, C.-M., Luo, R.: MEGAHIT: an ultra-fast single-node solution for large and complex metagenomics assembly via succinct de Bruijn graph. Bioinformatics 31(10), 1674–1676 (2015)

18. Li, H., Durbin, R.: Fast and accurate short read alignment with Burrows-Wheeler transform. Bioinformatics 25(14), 1754–1760 (2009)

19. Li, H., Handsaker, B., Wysoker, A.: The Sequence Alignment/Map format and SAMtools. Bioinformatics 25(16), 2078–2079 (2009)

20. Muggli, M.D., Bowe, A., Noyes, N.R.: Succinct colored de Bruijn graphs. Bioinformatics 33(20), 3181–3187 (2017)

21. Muir, P., Li, S., Lou, S.: The real cost of sequencing: scaling computation to keep pace with data generation. Genome Biology 17(1), 53 (2016)

22. Novak, A.M., Garrison, E., Paten, B.: A graph extension of the positional Burrows-Wheeler transform and its applications. Algorithms for Molecular Biology 12 (2017)

23. Ondov, B.D., Treangen, T.J., Melsted, P.: Mash: fast genome and metagenome distance estimation using MinHash. Genome Biology 17(1), 132 (2016)

24. Pandey, P., Bender, M.A., Johnson, R.: deBGR: an efficient and near-exact representation of the weighted de Bruijn graph. Bioinformatics 33(14), i133–i141 (2017)

25. Paten, B., Diekhans, M., Earl, D.: Cactus Graphs for Genome Comparisons. Journal of Computational Biology 18(3), 469–481 (2011)

26. Paten, B., Novak, A.M., Eizenga, J.M.: Genome graphs and the evolution of genome inference. Genome research 27(5), 665–676 (2017)

27. Pevzner, P.A., Tang, H., Waterman, M.S.: An Eulerian path approach to DNA fragment assembly. Proceedings of the National Academy of Sciences 98(17), 9748–9753 (2001)

28. Raman, R., Raman, V., Satti, S.R.: Succinct indexable dictionaries with applications to encoding k-ary trees, prefix sums and multisets. ACM Transactions on Algorithms 3(4), 43-es (2007)

29. Solomon, B., Kingsford, C.: Fast search of thousands of short-read sequencing experiments. Nature Biotechnology 34(3), 300–302 (2016)

30. Solomon, B., Kingsford, C.: Improved search of large transcriptomic sequencing databases using split sequence bloom trees. In: Lecture Notes in Computer Science (including subseries Lecture Notes in Artificial Intelligence and Lecture Notes in Bioinformatics), pp. 257–271 (2017)

31. Tischler, G., Leonard, S.: biobambam: tools for read pair collation based algorithms on BAM files. Source Code for Biology and Medicine 9(1), 13 (2014)

32. Turnbaugh, P.J., Ley, R.E., Hamady, M.: The human microbiome project: exploring the microbial part of ourselves in a changing world. Nature 449(7164), 804–810 (2007)

33. Walter, K., Min, J.L., Huang, J.: The UK10K project identifies rare variants in health and disease. Nature 526(7571), 82–90 (2015)

34. Zerbino, D.R., Birney, E.: Velvet: Algorithms for de novo short read assembly using de Bruijn graphs. Genome Research 18(5), 821–829 (2008)

35. Zhang, G., Li, C., Li, Q.: Comparative genomics reveals insights into avian genome evolution and adaptation. Science 346(6215), 1311–1320 (2014)

36. Zhang, Q., Pell, J., Canino-Koning, R.: These are not the k-mers you are looking for: efficient online k-mer counting using a probabilistic data structure. PloS one 9(7), e101271 (2014)

